# A Reproducible Cortical Model of Focal Epileptiform Activity with a Multi-Biomarker Analysis Pipeline

**DOI:** 10.64898/2026.10.06.756259

**Authors:** C López-Cabezón, T Gener, A Polo-Rodríguez, MV Puig

**Author notes:** **Correspondence**: Prof. M. Victoria Puig, PhD.

## Abstract

The development of emerging optical and epicortical neuromodulatory strategies requires experimental models in which focal pathological activity can be generated and monitored within an accessible cortical region. Here, we established an acute cortical model of focal epileptiform activity in anesthetized mice based on local microinjection of 4-aminopyridine (4-AP) into the barrel cortex. This preparation provides precise spatial and temporal control of the epileptiform focus while enabling simultaneous multichannel electrophysiological recording. To objectively characterize the induced activity, we implemented a Python-based analysis pipeline integrating complementary electrophysiological biomarkers, including frequency-specific power, multi-unit activity (MUA), pathological fast ripples (250–650 Hz), and epileptiform bursts. Local 4-AP administration produced marked increases in high-frequency (HF; 100–250 Hz) power, multi-unit spike rate, fast ripple rate, and cumulative burst duration. Pharmacological sensitivity of the framework was evaluated using the antiseizure drug carbamazepine, which consistently attenuated these 4-AP-induced electrophysiological alterations, bringing values close to those observed in vehicle-treated animals. Together, this study provides a reproducible experimental and analytical framework for investigating focal cortical epileptiform activity and for the future evaluation of pharmacological, optical, and electrical interventions targeting superficial cortical networks.

## INTRODUCTION

Epilepsy is one of the most prevalent neurological disorders, affecting over 70 million people worldwide and carrying an increased risk of premature mortality (Thijs et al., 2019). Although appropriate treatment can achieve seizure freedom in a substantial proportion of patients, approximately one-third remain resistant to currently available antiseizure medications, highlighting the continued need for experimental models that support mechanistic studies and the development of new therapeutic strategies. Focal seizures originate within a localized brain region and represent the predominant seizure type in a large proportion of patients. Understanding how focal epileptiform activity emerges, evolves, and responds to therapeutic intervention therefore remains a major objective of preclinical epilepsy research.

A wide variety of experimental models has been developed to study seizure generation and the longer-term process of epileptogenesis, each reproducing different aspects of the human disorder (Avoli et al., 2016; Löscher, 2017). Chemoconvulsant models based on kainate or pilocarpine have been particularly valuable for investigating temporal lobe epilepsy, long-term epileptogenesis, and spontaneous recurrent seizures, although they may also involve substantial variability in seizure severity, mortality, and neuropathological outcome (Arida et al., 1999; Hamani and Mello, 2002; Lévesque and Avoli, 2013; Kandratavicius et al., 2014; Twele et al., 2017; Jiang and Wang, 2025). Intrahippocampal kainate, in particular, provides a well-established model of chronic temporal lobe epilepsy, but its deep anatomical location and long-term phenotype, together with the spontaneous and temporally unpredictable nature of seizure occurrence, make it less suitable for experimental paradigms requiring direct access to a superficial focal site and precise control over when epileptiform activity emerges during a recording session.

Our primary objective was to establish an acute cortical model of focal epileptiform activity suitable for simultaneous electrophysiological characterization and testing of surface-based therapeutic interventions. This is particularly relevant given the emergence of light-controlled pharmacological strategies and epicortical interfaces capable of electrophysiological monitoring and focal stimulation (Camerin et al., 2024; Prokop et al., 2026). Experiments were performed under isoflurane anesthesia to provide a stable preparation for focal microinjection and multichannel recordings. This experimental context also influenced the choice of chemoconvulsant: isoflurane enhances GABA_A_ receptor-mediated inhibition and reduces neuronal excitability (Jia et al., 2008), and anesthesia has been shown to modify the outcome of intrahippocampal kainate models (Twele et al., 2016). In contrast, focal cortical administration of 4-aminopyridine (4-AP) reliably induces epileptiform activity under isoflurane anesthesia (Zhao et al., 2015; Nagappan et al., 2019; Yang et al., 2021).

A further challenge in experimental epilepsy concerns the objective identification and quantification of epileptiform activity. Conventional seizure analysis frequently relies on visual inspection of electrophysiological recordings, which is subjective, time-consuming, and associated with substantial inter-rater variability (Jing et al., 2023; Aanestad et al., 2024; Kamintsky et al., 2025). This has encouraged the development of automated and multiparametric approaches that capture complementary features of pathological neural activity (Zuev et al., 2025; Wang et al., 2026). Changes in spectral composition and neuronal firing represent distinct aspects of network hyperexcitability and may respond differently to therapeutic intervention. Their integration within a common analytical framework may therefore provide a more informative and reproducible characterization of focal epileptiform activity than reliance on a single electrophysiological endpoint.

Here, we established an acute cortical model of focal epileptiform activity in anesthetized mice based on local 4-AP microinjection into the barrel cortex, combined with multichannel electrophysiological recording and a Python-based multi-biomarker analysis pipeline. The S1 barrel cortex was selected for its superficial location and well-defined barrel and laminar organization, and because it has previously been used to combine focal 4-AP-induced epileptiform activity with optogenetic modulation, supporting its suitability for experimental studies of focal cortical epilepsy (Assaf and Schiller, 2016). The pipeline was implemented to quantify complementary biomarkers including oscillatory power, MUA, pathological fast ripples, and epileptiform bursts. Pharmacological sensitivity of the framework was assessed using carbamazepine (CBZ), a widely used antiseizure medication whose anticonvulsant action is primarily associated with inhibition of voltage-gated sodium channels (Kuo et al., 1997), thereby limiting direct mechanistic overlap with the predominantly GABA_A_ receptor-mediated effects of isoflurane (Jia et al., 2008). Together, this experimental and analytical framework provides a reproducible platform for investigating focal cortical epileptiform activity and for evaluating promising pharmacological interventions while supporting future studies of optical and electrical epicortical neuromodulatory strategies.

## MATERIALS AND EQUIPMENT

### Animals and Housing

Young adult male and female C57BL/6J mice (8–12 weeks old; 19–27 g) were obtained from Janvier Labs. This age range was selected based on preliminary observations indicating more consistent electrophysiological responses to 4-AP, thereby improving the reproducibility of the model. Animals were housed under controlled environmental conditions at 22 ± 2 °C and 60% relative humidity, with a 12:12 h light–dark cycle (lights off from 8:00 p.m. to 8:00 a.m.). Experimental procedures were performed between 9:00 a.m. and 5:00 p.m., during the light phase of the cycle. Standard pellet food and water were available *ad libitum* until the experimental session. Animals were euthanized at the end of the experiment.

### Drugs and Reagents

4-Aminopyridine (4-AP; A78403, Sigma-Aldrich) was used to induce focal cortical epileptiform activity. A 30 mM stock solution of 4-AP was prepared in phosphate-buffered saline (PBS; 524650, Merck Millipore) and stored at −20 °C until use. Carbamazepine (CBZ; C4024, Sigma-Aldrich) was used for pharmacological validation of the model. CBZ was initially dissolved in DMSO and subsequently diluted in PBS to prepare a 2 mM stock solution containing 6% (v/v) DMSO, which was stored at −20 °C until use.

On the day of each experiment, working solutions were freshly prepared in PBS. For the 4-AP group, the 30 mM stock solution was diluted to a final concentration of 15 mM, an intermediate concentration within the range previously reported for focal cortical 4-AP models (5–25 mM) (Zhao et al., 2015; Nagappan et al., 2019; Yang et al., 2021), and DMSO was added to obtain a final concentration of 3% (v/v). For co-treatment experiments, 4-AP and CBZ were combined in the same working solution to obtain final concentrations of 15 mM 4-AP together with either 600 μM or 1 mM CBZ, with the final DMSO concentration adjusted to 3% (v/v). Vehicle-control animals received PBS containing 3% DMSO but no 4-AP or CBZ. All working solutions were maintained at 4 °C until loading into the injection syringe and discarded after use.

Atropine sulfate (3545024, Braun Medical) was administered subcutaneously at 0.05 mg/kg before surgery to reduce airway secretions during anesthesia. Isoflurane (3369 ESP, Isoflurin, Fatro) was used for induction and maintenance of general anesthesia.

For histological verification, five animals selected for tissue processing were transcardially perfused with PBS followed by 4% paraformaldehyde (PFA; J61899.AP, Thermo Scientific Chemicals). Hoechst 33342 (ref. H3570, Invitrogen) was used as a nuclear counterstain at a 1:10,000 dilution in PBS, and sections were mounted using Entellan mounting medium (ref. 1.07960.0500, Sigma-Aldrich).

### Recording Electrodes

Electrophysiological recordings were initially performed using tungsten stereotrodes during the development and optimization of the experimental model. Once the recording conditions and model parameters had been established, additional animals from each experimental group were recorded using 16-channel silicon probes (NeuroNexus Technologies, Inc., USA) to provide laminar recordings across the S1BF. Tungsten stereotrodes consisted of two twisted strands of 25 μm tungsten wire (Advent Research Materials, UK), whereas the 16-channel silicon probes (A1x16-5mm-50-177-A16; NeuroNexus) were connected through an Adpt-A16-OM32 adapter and comprised vertically aligned recording sites spaced 50 μm apart along a single shank, spanning a total distance of 750 μm (Figure 1B,C). Each silicon probe was reused for recordings in 3–4 different animals. Between experiments, probes were immersed for 10 min in a catalase-containing enzymatic cleaning solution (Arion Cronos, Disop) to remove biological residues, followed by immersion in 70% ethanol.

**FIGURE 1:**
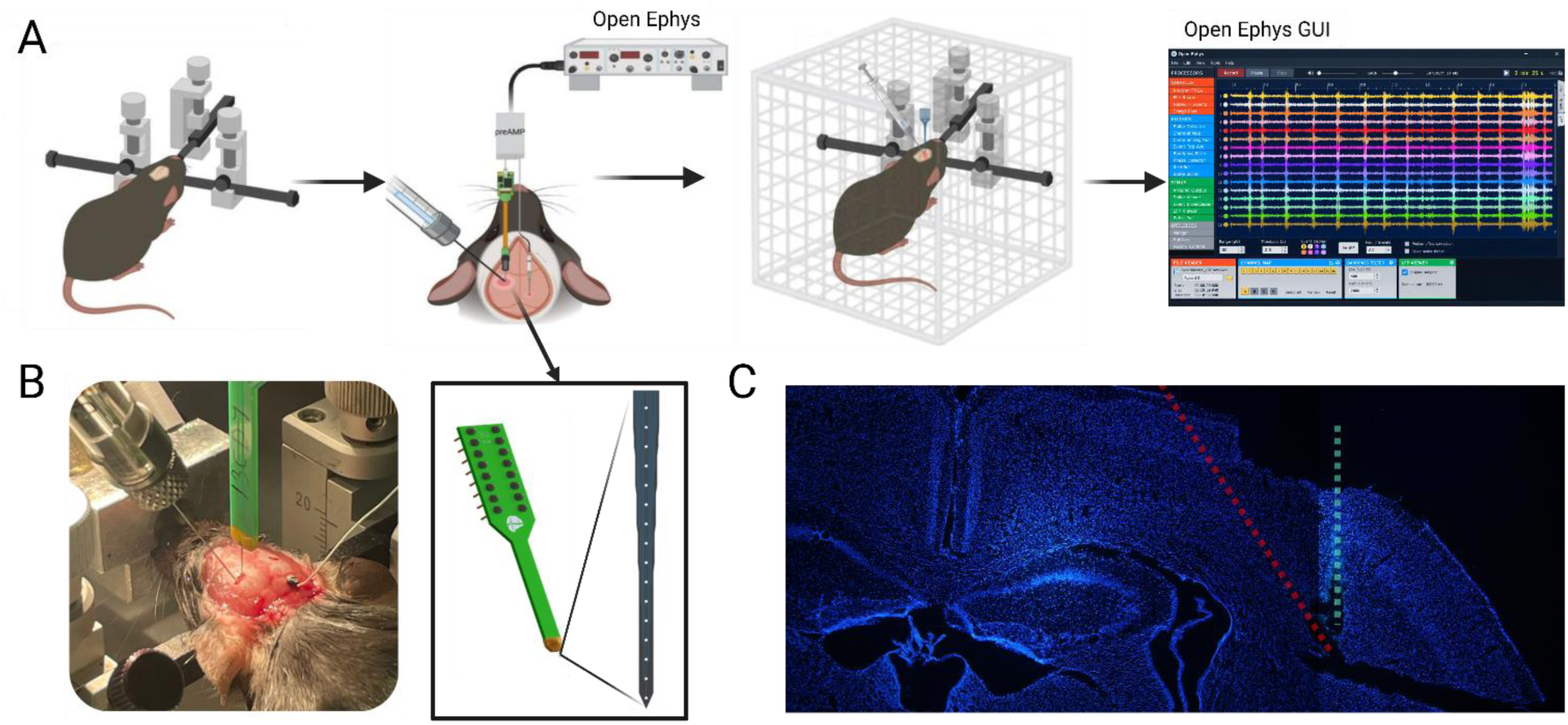
Surgical setup, microinjection, and electrophysiological recording configuration in the barrel cortex. **(A)** Schematic of the experimental pipeline: stereotaxic frame fixation, probe and syringe implantation, placement in a Faraday cage for signal shielding, and real-time multichannel signal acquisition using Open Ephys software. **(B)** Representative photograph showing the in vivo surgical site setup with stereotaxic micromanipulators positioning the injection capillary needle, recording probe, and ground wire. Design layout of the linear silicon probe array. **(C)** Anatomical localization of the region of interest (ROI) in the primary somatosensory cortex, barrel field (S1BF), showing histological trace verification alongside stereotaxic brain atlas coordinates.

## METHODS

### Surgical Procedures

All procedures were performed under general anesthesia with inhaled isoflurane (3–4% for induction and during surgery, followed by 1% for maintenance throughout the remainder of the experimental session). To reduce airway secretions, atropine (0.05 mg/kg) was administered subcutaneously 15 min before the start of surgery. A small craniotomy was made over the barrel field of the primary somatosensory cortex (S1BF; AP: −1.22 mm; ML: ±3.0 mm relative to bregma) to provide access for both electrophysiological recording and drug administration. The right or left hemisphere was selected at random for each experiment. A dual-arm stereotaxic frame was used to independently position the recording probe, which was first lowered into the target region, and a Hamilton syringe (75RN, 5 μL; Hamilton Company), subsequently inserted 650 μm at a 30° angle for drug delivery. A single microscrew was placed over the cerebellum and used as the common ground electrode (Figure 1B).

### Electrophysiological Recordings

Recording electrodes were positioned in the S1BF. Tungsten stereotrodes were lowered to DV −0.6 mm, targeting approximately the layer IV/V boundary, whereas the tip of the 16-channel silicon probes was lowered to DV −0.9 mm. The recording sites, distributed over a 750-μm span along the probe shank, enabled simultaneous sampling across multiple cortical layers of the S1BF (Figure 1B,C). Neural activity was continuously monitored during electrode advancement to assist positioning within the target cortical region. Electrode impedance was measured before each recording and ranged from 100 to 400 kΩ for tungsten stereotrodes and from 0.7 to 1.4 MΩ for the 16-channel silicon probes. During electrophysiological acquisition, animals were positioned inside a custom-built aluminum Faraday cage (40 × 50 × 55 cm). Signals were acquired using the Open Ephys system (version 0.4.3.3) with Intan RHD2132 amplifier boards, sampled at 30 kHz, and band-pass filtered between 0.1 and 6,000 Hz. Continuous recordings were stored as consecutive 10-min files in .kwik format.

### Drug Delivery and Experimental Groups

Following placement of the recording electrode, the Hamilton syringe was preloaded with the corresponding drug or vehicle solution and positioned approximately 1 mm lateral to the cortical entry point of the recording electrode. The syringe was then inserted 650 μm into the S1BF at a 30° angle. This configuration allowed local microinjection adjacent to the recorded area while minimizing interference with electrode placement. After needle placement, the Faraday cage was closed and a 10-min baseline period was recorded before drug administration (Figure 1A, Figure 2A). Solutions were then delivered at a rate of 3 μL/min, corresponding to a total volume of 1 μL over 20 s, using a microinjector (53313, Stoelting Co.). Neural activity was subsequently recorded continuously for 60 min after injection to characterize the evoked response. S1BF activity was assessed under four experimental conditions: vehicle control (PBS + DMSO), 4-AP alone (15 mM), and co-administration of 4-AP with carbamazepine at 600 μM and 1 mM concentrations. All animals were euthanized at the end of the recording session (Figure 2A,B).

**FIGURE 2:**
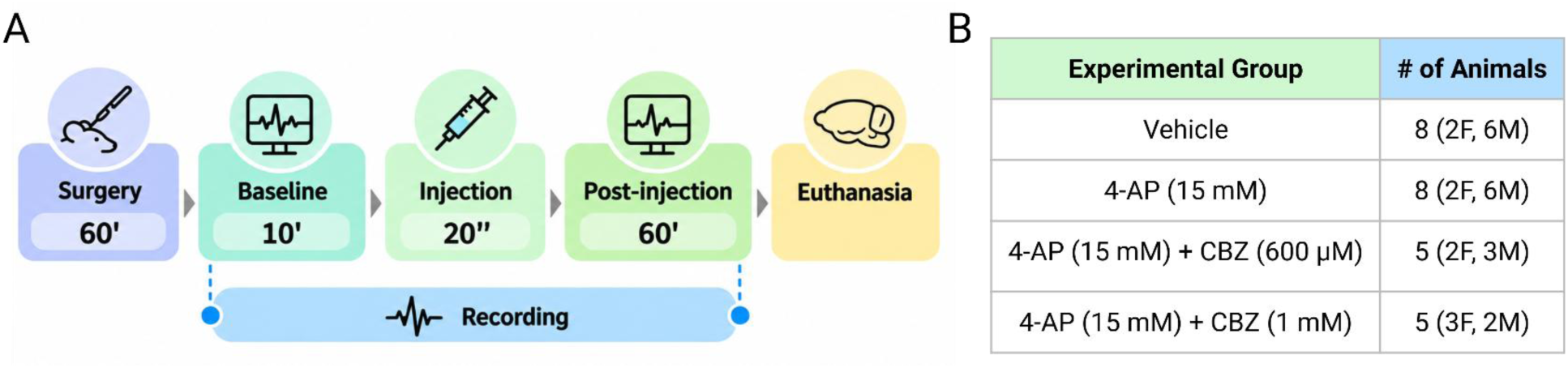
Timeline and experimental groups of the study. **(A)** Schematic representation of the experimental workflow and time course. Following a 60-min surgical preparation period, continuous electrophysiological recording comprised a 10-min baseline period, a 20-s injection period, and a 60-min post-injection observation period, after which animals were euthanized. (**B**) Summary of the experimental groups, including animal allocation by sex (F, females; M, males). The vehicle control group included 3 animals recorded with stereotrodes and 5 with silicon probes; the 4-AP group (15 mM) included 6 animals recorded with stereotrodes and 2 with silicon probes; the 4-AP + CBZ 600 μM group included 1 animal recorded with a stereotrode and 4 with silicon probes; and the 4-AP + CBZ 1 mM group included 3 animals recorded with stereotrodes and 2 with silicon probes.

### Histology

A subset of animals was subsequently transcardially perfused, and the brains were collected and sectioned for histological verification of the placement of the recording probe and Hamilton syringe. Sections were stained with Hoechst 33342, a fluorescent nuclear counterstain that labels cell nuclei irrespective of cell type, allowing visualization of general tissue architecture and confirmation of the electrode and syringe tracks within the S1BF in representative samples (Figure 1C). Sections were washed 3 times for 10 min with PBS under agitation and then incubated for 10 min with Hoechst solution at RT. The washing steps were then repeated (3 × 10 min), and sections were mounted onto glass slides. Once mounted, the sections were coverslipped with Entellan mounting medium. The slides were then left to dry before imaging on an Olympus BX51 fluorescence microscope equipped with an Olympus DP74 camera, using cellSens Dimension software (version 4.2.1).

### Electrophysiological Analyses

Custom Python scripts were executed in the Spyder environment to quantify electrophysiological biomarkers of focal epileptiform activity. Prior to analysis, raw signals stored as consecutive 10-min .kwik files were imported and converted to .mat format. The individual 10-min segments from each channel were then concatenated in chronological order to generate a continuous 70-min recording, from which raw and z-score-normalized spectrograms, multi-unit activity (MUA), and fast ripples (250–650 Hz) were obtained (Figure 3, Figure 4).

**FIGURE 3:**
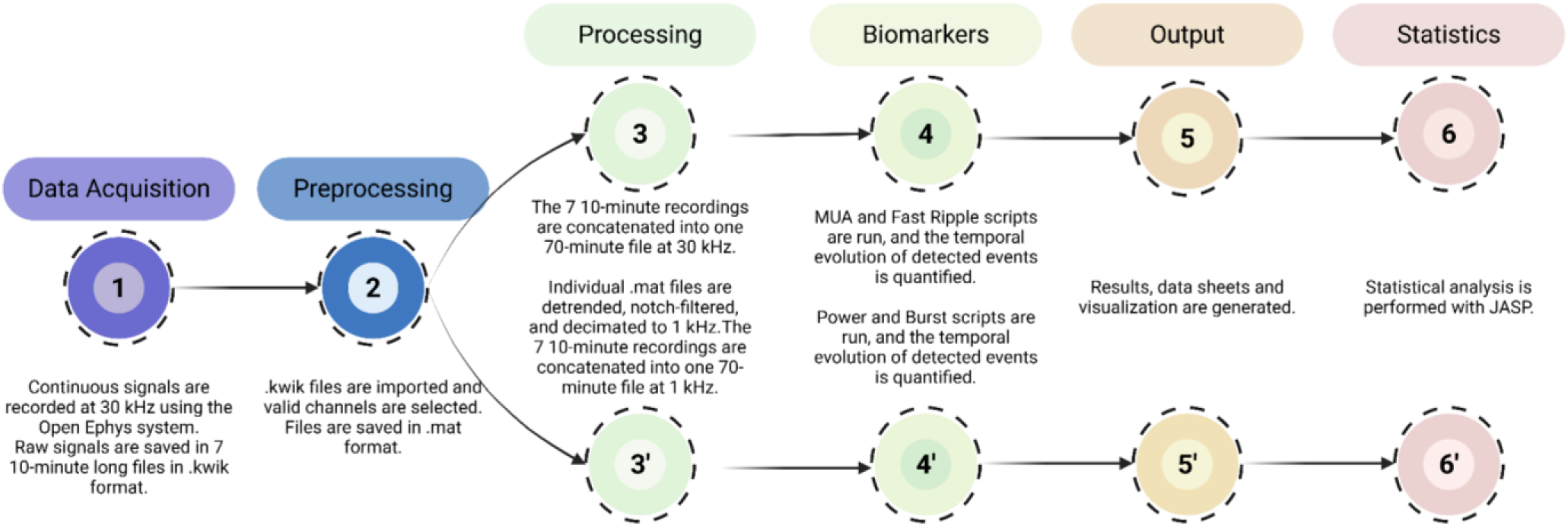
Signal processing and feature extraction pipeline for electrophysiological recordings. Sequential workflow for preprocessing raw Open Ephys .kwik data, feature extraction and biomarker detection, followed by generation of analysis outputs for statistical testing in JASP. The custom Python-based analysis code used in this study is available through the laboratory’s GitHub repository.

**FIGURE 4:**
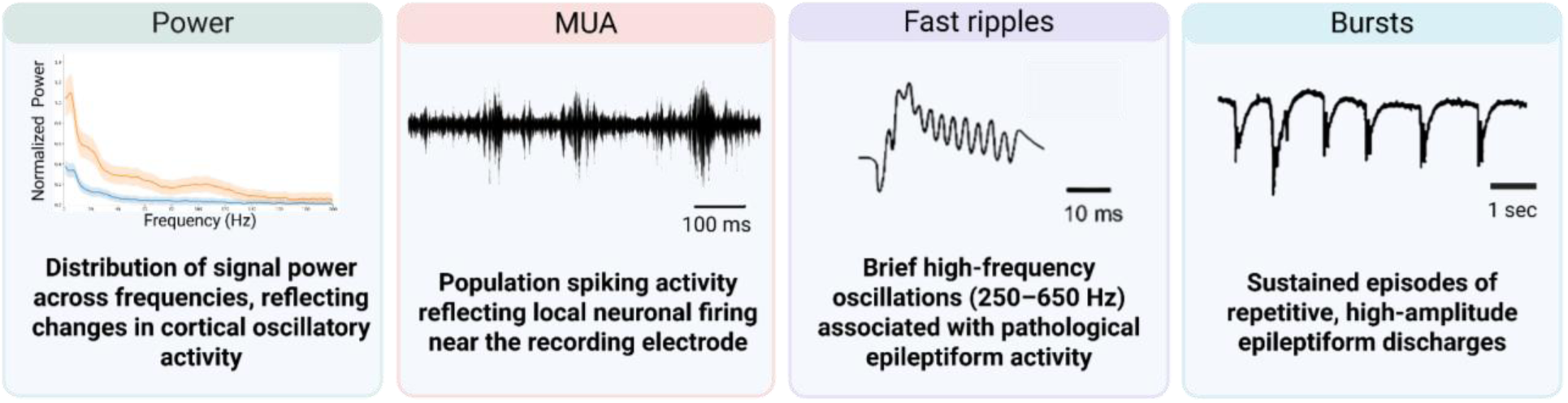
Electrophysiological features used to characterize focal epileptiform activity. Representative schematic examples of the principal electrophysiological alterations assessed in the analysis pipeline: changes in spectral power distribution, changes in multi-unit spiking activity (MUA), pathological fast ripple oscillations (250–650 Hz), and sustained epileptiform burst discharges.

In parallel, signals from each recording .mat channel were detrended to remove slow baseline drifts, notch-filtered at 50 and 100 Hz to minimize power-line interferences, downsampled from 30 kHz to 1 kHz (decimation factor of 30) and finally concatenated in a 70-min recording. These recordings were utilized for power quantification, and burst detection (Figure 3, Figure 4). Detailed detection and quantification procedures for each electrophysiological biomarker are described in the corresponding subsections below.

#### Oscillatory power

Spectrograms were computed for each channel directly from the original 30-kHz recordings using a short-time Fourier transform with a 40-s window and were visualized over the 0–650 Hz frequency range. Power was expressed in decibels (10 log₁₀) for visualization. Spectrograms were generated over the full 70-min recording, with the first 10 min defined as baseline. In addition to the raw power spectrograms, baseline-normalized spectrograms were generated by normalizing power independently at each frequency bin relative to its distribution during the 10-min baseline period. Specifically, power values were centered to the corresponding baseline median and scaled by the baseline median absolute deviation (MAD), providing a robust z-score-like normalization of time-resolved spectral changes. The resulting baseline-normalized spectrograms were smoothed along the frequency axis using a Gaussian filter for visualization (Figure 5).

**FIGURE 5:**
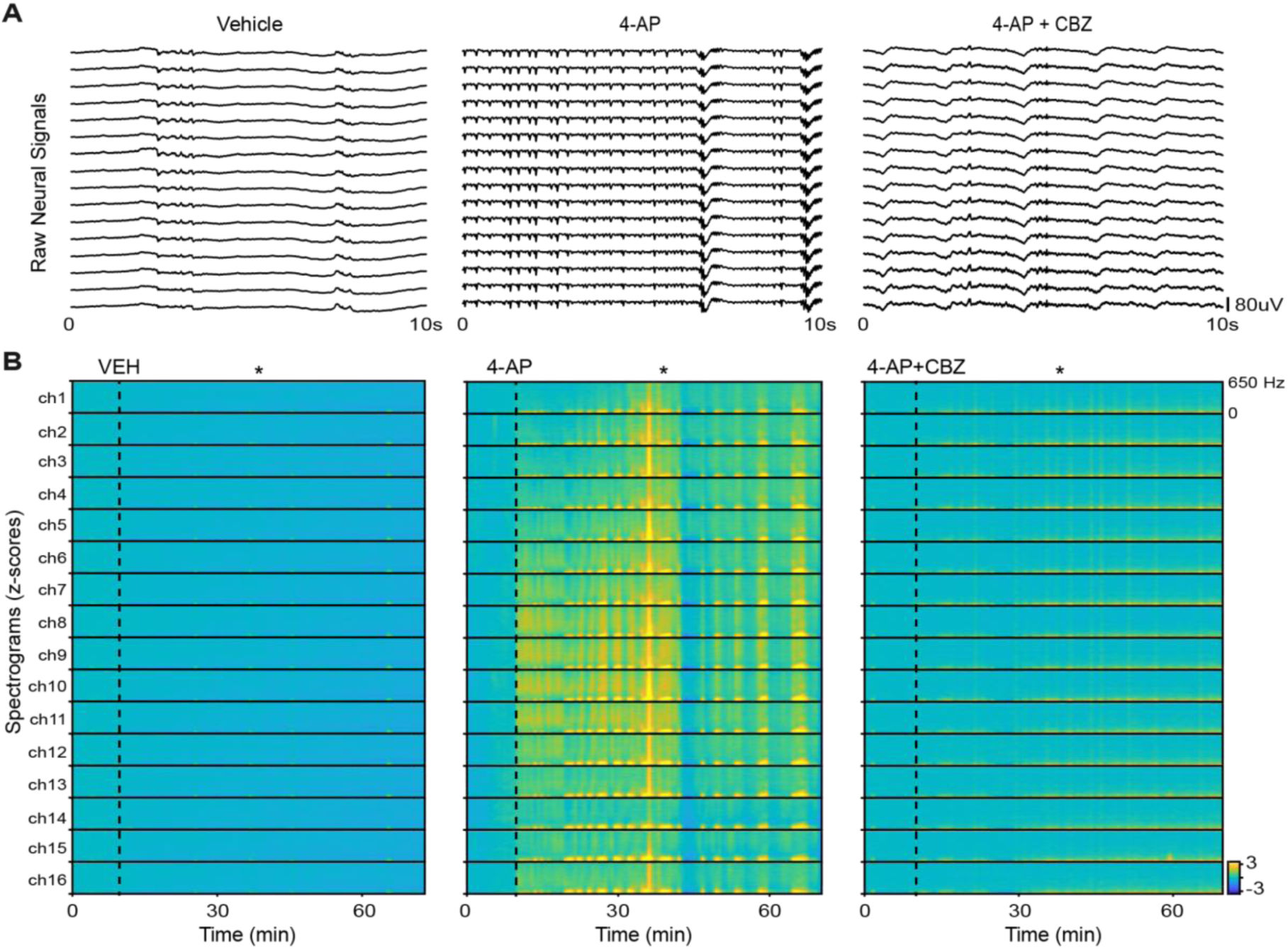
Representative electrophysiological and spectral responses following 4-AP administration and CBZ co-treatment. **(A)** Representative 10-s electrophysiological traces recorded with 16-channel silicon probes in vehicle, 4-AP (15 mM), and 4-AP (15 mM) + CBZ (1 mM) groups approximately 30 min after microinjection. **(B)** Baseline-normalized spectrograms (0–650 Hz) from representative animals in the same experimental groups, displayed across the 16 recording sites over the complete 70-min recording period. The first 10 min correspond to baseline, and the vertical dashed line indicates the time of microinjection. Asterisks indicate the approximate time window from which the representative traces shown in panel A were extracted (∼30 min after injection). Warmer colors indicate increases in spectral power relative to baseline, whereas cooler colors indicate decreases.

For quantitative spectral analyses, the signal was downsampled to 1 kHz, and power was quantified separately in the theta (4–12 Hz), alpha (13–18 Hz), beta (18–30 Hz), gamma (30–100 Hz), and high-frequency (HF; 100–250 Hz) bands. For each channel, band-specific power was calculated independently and averaged within consecutive 10-min intervals. Values were normalized to the corresponding baseline power measured during the first 10 min of the recording, yielding a baseline ratio of 1 for each animal. Because baseline values were fixed by normalization, only the six post-injection intervals were included in the statistical analyses to characterize the temporal evolution of oscillatory activity following vehicle, 4-AP, or 4-AP + CBZ administration.

#### Multi-unit spiking activity (MUA)

Continuous signals were band-pass filtered between 300 and 3,000 Hz using a fourth-order zero-phase Butterworth filter to isolate MUA. For each channel, the detection threshold was first established by visual inspection using Offline Sorter v4 (Plexon Inc.), following the criterion described in (Delgado-Sallent et al., 2023). Thresholds typically corresponded to approximately −2.5σ, although the exact multiplier (*k*) was adjusted individually for each channel. Spike detection was subsequently performed in Python using a negative threshold of −*k* × σ, with σ robustly estimated for each channel as 1.4826 times the median absolute deviation (MAD) of the filtered signal (Quiroga et al., 2004). A refractory period of 1.5 ms was imposed to prevent multiple detections of the same event. Spike rates were quantified in consecutive 5-min blocks and expressed as spikes per minute (spikes/min), with block duration taken into account for incomplete terminal segments (Figure 4). For statistical analyses, consecutive 5-min blocks were averaged to obtain values for each 10-min interval.

#### Fast ripples

Fast ripples are considered a pathological form of high-frequency oscillation and are strongly associated with epileptogenic tissue (Bragin et al., 1999; Staba et al., 2002; Weiss et al., 2020). Fast ripples were detected from signals retained at the original 30-kHz sampling rate. Signals were band-pass filtered between 250 and 650 Hz, and the instantaneous amplitude envelope was obtained using the Hilbert transform. Detection followed an envelope-based approach conceptually related to that described by Weiss and colleagues (Weiss et al., 2020), in which oscillatory events are identified from the normalized signal envelope and subsequently screened using temporal, amplitude, and spectral criteria. For each channel, the detection threshold was defined as the median plus four times the median absolute deviation (MAD) of the envelope calculated over the entire recording and was applied consistently to both baseline and post-injection periods. Candidate events were required to have a minimum duration of 3 ms. Events were excluded when peak-to-peak amplitude exceeded 500 µV, absolute waveform skewness exceeded 2.5, or less than 70% of their spectral power, estimated using Welch’s method, was contained within the 250–650 Hz band. These criteria were applied to minimize contamination by large-amplitude artifacts and broadband transient activity (Figure 4). Fast ripple counts were expressed as events per minute and quantified over consecutive 10-min intervals.

#### Epileptiform Bursts

Epileptiform bursts were defined as sustained periods of increased pathological network activity characterized by repetitive waveform deflections and enhanced spectral power relative to baseline (Figure 4). Burst detection was performed on the downsampled (1 kHz) LFP signal using a composite spectral–morphological approach implemented in the present analysis pipeline and inspired by previous quantitative approaches to epileptiform burst characterization (Heining et al., 2019). The continuous signal from each channel was divided into 256-ms windows with 50% overlap, and four complementary features were extracted from each window: line length, reflecting the cumulative amplitude variation of the signal within each analysis window and therefore increasing with larger and/or more frequent waveform deflections (Esteller et al., 2001); spectral power in the 4–40 Hz range (E₄–₄₀ Hz); spectral power in the 80–200 Hz range (E₈₀–₂₀₀ Hz); and spectral entropy (SE), reflecting the distribution of power across frequencies (Inouye et al., 1991). Band power was estimated using multitaper spectral analysis.

Each feature was converted to a robust z-score relative to the 10-min baseline period using the corresponding baseline median and median absolute deviation (MAD), scaled by 1.4826. The normalized features were then combined into an integrated index of burst activity, calculated as:

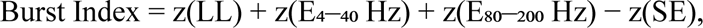

where z denotes baseline-normalized feature values. Thus, increases in line length and spectral power, together with a reduction in spectral entropy, resulted in higher index values and indicated sustained bursting activity.

Bursts were identified by applying separate onset and offset thresholds to the composite index. Event onset was defined when the index exceeded an upper threshold of 2.8, and the event was considered to end when it subsequently fell below a lower threshold of 1.6. The use of distinct onset and offset thresholds minimized event fragmentation caused by brief fluctuations around the detection threshold. Only events lasting at least 7 s were classified as epileptiform bursts. The total time spent in burst activity was then summed within consecutive 10-min blocks to characterize the temporal evolution of epileptiform activity throughout the recording.

#### Statistical analyses

All statistical analyses were performed using JASP open-source software (version 0.19.3). Electrophysiological biomarkers, including band power, spikes per minute, fast ripple rate, and cumulative burst duration, were analyzed using mixed-design ANOVAs, with Time as the within-subjects factor and Treatment as the between-subjects factor. For normalized band power and MUA, Time comprised six consecutive 10-min post-injection intervals, as baseline values were fixed at 1 by normalization and were therefore excluded from the statistical analyses. For fast ripple rate and cumulative burst duration, Time comprised seven levels: the 10-min baseline period and six consecutive 10-min post-injection intervals.

Primary statistical analyses compared the vehicle, 4-AP, and 4-AP + CBZ 1 mM groups. The lower CBZ concentration (600 μM) was analyzed separately in Supplementary Figure S1. Sphericity was assessed using Mauchly’s test, with Greenhouse–Geisser correction applied when the assumption was violated. To further characterize the temporal profile of treatment effects, pairwise comparisons between experimental groups were performed at individual post-injection intervals following the mixed-design ANOVA, with Holm correction for multiple comparisons. Statistical significance was set at p ≤ 0.05. Data are reported as mean ± SEM unless otherwise indicated.

## RESULTS

In an initial set of experiments, 4-AP was co-administered with CBZ at 600 µM (n = 5 mice; Figure 2). Responses across electrophysiological biomarkers showed only a partial rescue of 4-AP-associated effects at this concentration (Supplementary Figure S1). Therefore, a higher CBZ concentration (1 mM) was evaluated in a separate group and is presented in the main analyses. Accordingly, the primary statistical analyses focused on three experimental groups: vehicle (n = 8), 4-AP 15 mM (n = 8), and 4-AP 15 mM + CBZ 1 mM (n = 5; Figure 2).

Representative 10-s electrophysiological traces obtained with the 16-channel silicon probes are shown for the vehicle, 4-AP, and 4-AP + CBZ groups around minute 30 following microinjection (Figure 5A). Compared with vehicle, 4-AP induced marked changes in the amplitude and temporal organization of cortical activity, whereas these alterations appeared greatly attenuated following CBZ co-administration. To visualize the temporal and spectral evolution of these responses across cortical depth, baseline-normalized spectrograms were generated for each recording site over the full 70-min recording and across the 0–650 Hz frequency range (Figure 5B). Following 4-AP administration, spectral power increased markedly across frequencies below 100 Hz along the electrode track, with the strongest changes at intermediate recording sites and comparatively weaker effects at more superficial sites. This depth-dependent pattern was markedly reduced following CBZ co-administration. These spectrograms are intended to illustrate the characteristic spectral patterns observed within each group and were not used for statistical analysis. Representative animals were selected based on their correspondence with the predominant response observed within each experimental group.

### 4-AP induces rapid and persistent increases in high-frequency power that are attenuated by CBZ co-administration

Quantitative spectral analyses were performed separately for the theta (4–12 Hz), alpha (13–18 Hz), beta (18–30 Hz), gamma (30–100 Hz), and high-frequency (HF; 100–250 Hz) bands. Power was normalized to each animal’s baseline activity (ratio = 1). Following 4-AP administration, HF power showed a strong and persistent increase, remaining elevated throughout the post-injection period. Gamma power also increased, although the response was less stable over time. Power in the theta, alpha, and beta bands was likewise elevated following 4-AP administration, but these changes were considerably less consistent across animals. Overall, HF power emerged as the most robust spectral biomarker of 4-AP-induced epileptiform activity in this model.

For HF power, mixed-design ANOVA revealed a significant main effect of Treatment (F₂,₁₈ = 18.12, p < 0.001). CBZ co-administration markedly suppressed the 4-AP-induced increase in HF power, with values remaining comparable to those observed in vehicle-treated animals throughout the post-injection period. Holm-corrected post hoc comparisons confirmed significantly higher HF power in the 4-AP group than in the 4-AP + CBZ group at all post-injection time points (Figure 6A). These findings identify HF power as a robust spectral measure of 4-AP-induced epileptiform activity and its pharmacological modulation. At the lower CBZ concentration (600 μM), this attenuation was only partial and showed greater inter-animal variability (Supplementary Figure S1).

**FIGURE 6:**
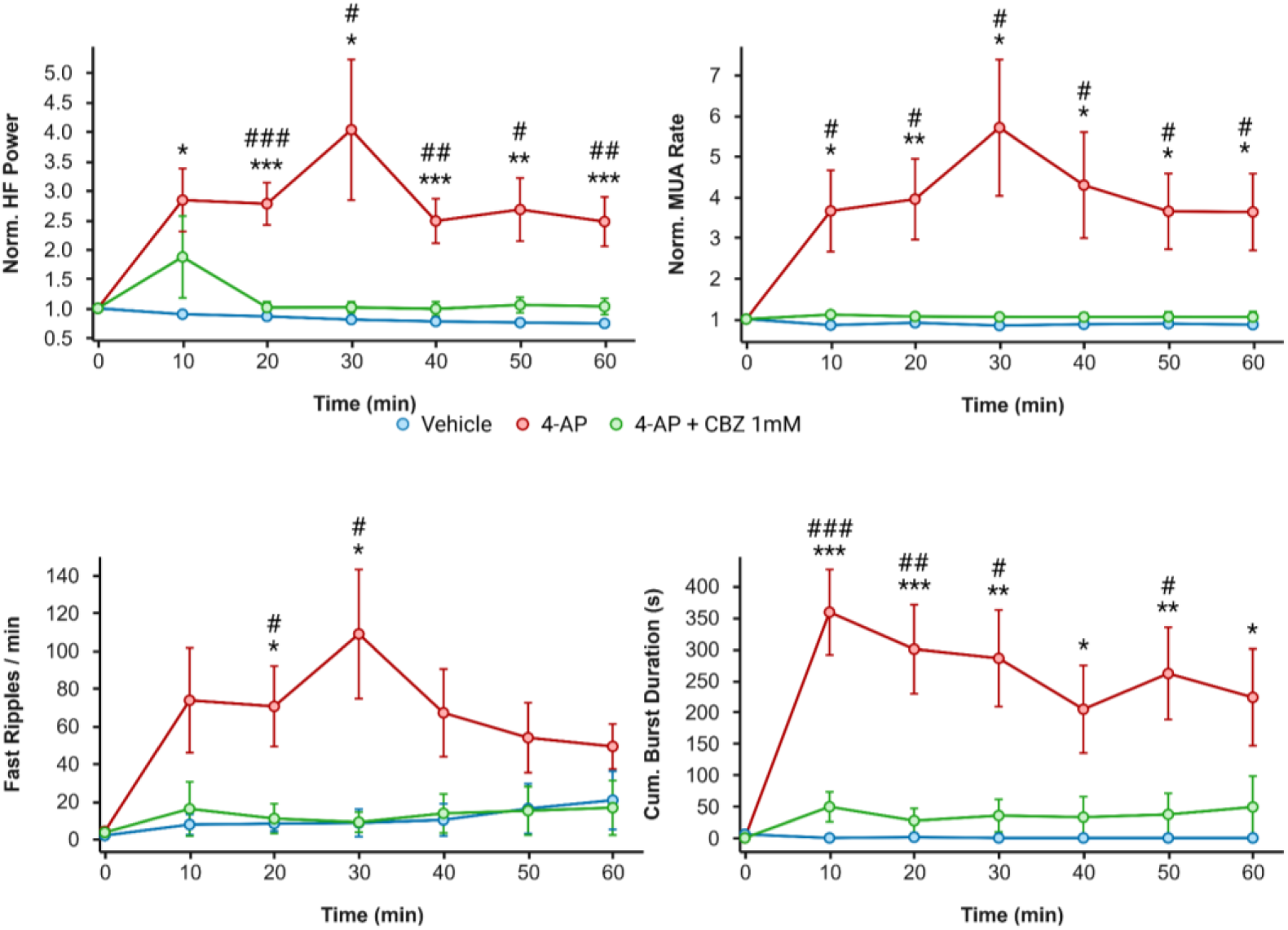
Temporal dynamics of electrophysiological biomarkers following 4-AP administration and CBZ co-treatment. Time courses of (**A**) normalized high-frequency (HF) power, (**B**) normalized multi-unit activity (MUA), (**C**) fast ripple rate, and (**D**) cumulative burst duration in vehicle, 4-AP (15 mM), and 4-AP (15 mM) + CBZ (1 mM) groups. HF power and MUA were normalized to each animal’s baseline value (baseline ratio = 1). Baseline is represented at 0 min, followed by six consecutive 10-min post-injection intervals. Data are presented as mean ± SEM. Pairwise comparisons between experimental groups were performed following the mixed-design ANOVA with Holm correction for multiple comparisons. Asterisks indicate significant differences between vehicle and 4-AP (*p < 0.05, **p < 0.01, ***p < 0.001), whereas hash symbols indicate significant differences between 4-AP and 4-AP + CBZ (#p < 0.05, ##p < 0.01, ###p < 0.001).

For gamma power, mixed-design ANOVA revealed a significant main effect of Treatment (F₂,₁₇ = 4.721, p = 0.023). CBZ co-administration also reduced the 4-AP-induced increase in gamma power, with values remaining comparable to those observed in vehicle-treated animals throughout the post-injection period. Holm-corrected post hoc comparisons showed significantly higher gamma power in the 4-AP group than in the 4-AP + CBZ group at 20, 30, and 40 min post-injection. Thus, gamma power was also sensitive to CBZ co-treatment, although the effect was less consistent over time than that observed for HF power.

### Normalized multi-unit activity rate is increased by 4-AP and restored to vehicle-comparable levels by CBZ co-administration

Multi-unit spike rate was normalized to each animal’s baseline activity (ratio = 1) to account for inter-individual differences in spontaneous firing under anesthesia. Analysis of the six post-injection intervals revealed a significant main effect of Treatment (mixed-design ANOVA; F₂,₁₈ = 7.814, p = 0.004), with no significant Time × Treatment interaction. To further characterize the temporal evolution of MUA, normalized activity was examined across successive 10-min post-injection intervals. The 4-AP group showed a progressive increase, reaching a maximum at 30–40 min post-injection (mean ratio = 5.701 ± 1.674), whereas vehicle-treated and 4-AP + CBZ animals remained close to baseline throughout the recording period. Holm-corrected post hoc comparisons showed significantly higher normalized MUA in the 4-AP group than in the 4-AP + CBZ group at all post-injection intervals, whereas no significant differences were detected between the 4-AP + CBZ and vehicle groups at any time point (Figure 6B). At 600 μM, CBZ produced only a partial reduction in the 4-AP-induced increase in MUA, with responses varying considerably across animals (Supplementary Figure S1).

### 4-AP-induced fast ripple activity is restored to vehicle-comparable levels by CBZ co-administration

Fast ripple activity was minimal during baseline and did not differ between experimental groups. Analysis across baseline and the six post-injection intervals revealed a significant main effect of Treatment (mixed-design ANOVA; F₂,₁₈ = 6.531, p = 0.007), whereas the Time × Treatment interaction did not reach statistical significance (p = 0.077). Following 4-AP administration, fast ripple activity increased progressively, reaching a maximum approximately 30 min after injection, and subsequently declined. In contrast, fast ripple activity remained low in vehicle-treated animals and was markedly attenuated by CBZ co-administration.

Holm-corrected post hoc comparisons showed significantly higher fast ripple activity in the 4-AP group than in the 4-AP + CBZ group at 20 and 30 minutes post-injection, whereas no significant differences were detected at the final three post-injection intervals. No significant differences were observed between the 4-AP + CBZ and vehicle groups at any time point, consistent with restoration to vehicle-comparable levels (Figure 6C). The lower CBZ concentration (600 μM) only partially attenuated the increase in fast ripple activity induced by 4-AP and showed marked inter-animal variability (Supplementary Figure S1).

### 4-AP increases cumulative burst duration that is attenuated by CBZ co-administration

Burst activity was negligible during baseline and did not differ between experimental groups. Cumulative burst duration was defined as the total time occupied by detected burst episodes within each 10-min interval. Analysis across baseline and the six post-injection intervals revealed significant main effects of Time and Treatment, together with a significant Time × Treatment interaction (mixed-design ANOVA; Time: F_2.68,48.30_ = 6.13, p = 0.002; Treatment: F_2,18_ = 12.08, p < 0.001; Time × Treatment: F_5.37,48.30_ = 5.28, p < 0.001), indicating that cumulative burst duration evolved differently across experimental groups following treatment.

Following 4-AP administration, cumulative burst duration increased rapidly, reaching a maximum at 10 min after injection before progressively declining (Figure 6D). In contrast, burst activity remained negligible in vehicle-treated animals throughout the recording period, whereas CBZ co-administration markedly attenuated the 4-AP-induced response. Holm-corrected post hoc comparisons showed significantly higher burst activity in the 4-AP group than in the 4-AP + CBZ group at 10, 20, 30, and 50 min post-injection. Burst activity in the 4-AP + CBZ group remained low, with no significant differences detected vs vehicle. At 600 μM, CBZ produced only a partial reduction in cumulative burst duration, with substantial variability between animals (Supplementary Figure S1).

## DISCUSSION

The present study establishes an acute model of focal cortical epileptiform activity based on local 4-AP administration in the mouse barrel cortex, combined with multichannel electrophysiological recordings and a Python-based multi-biomarker analysis pipeline. Local 4-AP administration produced marked and reproducible changes across several electrophysiological domains, including increased HF power, MUA, pathological fast ripple activity, and prolonged epileptiform burst episodes. Importantly, CBZ co-administration consistently attenuated the 4-AP-induced alterations across all principal electrophysiological readouts, with values approaching those observed in vehicle-treated animals, thereby providing pharmacological validation of the model.

The four principal electrophysiological readouts showed convergent but complementary responses to 4-AP and CBZ co-treatment. HF power and MUA exhibited particularly robust and sustained pharmacological separation between experimental groups, whereas fast ripple activity and cumulative burst duration captured complementary high-frequency oscillatory and episodic features of the epileptiform state. All four measures were markedly increased following 4-AP administration and substantially attenuated by CBZ co-treatment, with values approaching those observed in vehicle-treated animals. Taken together, the convergence of these complementary measures supports the use of a multi-biomarker strategy to characterize focal epileptiform activity and its pharmacological modulation. Because CBZ was administered simultaneously with 4-AP, however, these findings should be interpreted as evidence of sensitivity to anticonvulsant co-treatment rather than reversal of already established epileptiform activity.

Spectral analyses revealed that 4-AP-induced changes extended across multiple frequency bands, although their robustness differed substantially. HF power showed the strongest and most persistent response, remaining elevated throughout the post-injection period and providing clear pharmacological separation between the 4-AP and CBZ groups. Gamma power also increased following 4-AP administration and was sensitive to CBZ co-treatment, although this effect was less consistent over time. In contrast, changes in theta, alpha, and beta power were more variable across animals. These findings identify HF power as the most reliable spectral biomarker in this model and further support the value of combining spectral and non-spectral measures to characterize focal epileptiform activity.

A further advantage of the model is the superficial and spatially restricted location of the epileptiform focus. The barrel cortex provides a well-defined cortical target in which local pharmacological manipulation can be combined with multichannel electrophysiological recordings within a single experimental session. This configuration is particularly suitable for studies requiring direct access to the cortical surface, including emerging optical, electrical, and optoelectronic approaches. Photoswitchable carbamazepine analogues have recently demonstrated the feasibility of light-controlled pharmacological modulation in vivo (Camerin et al., 2024), while epicortical graphene-based neural interfaces enable broadband recording and focal electrical neuromodulation (Bonaccini-Calia et al., 2022; Prokop et al., 2026). The present preparation may therefore provide a controlled platform for evaluating how such interventions affect multiple electrophysiological features of focal cortical epileptiform activity.

Several limitations should nevertheless be considered. First, this is an acute preparation and does not reproduce the spontaneous recurrent seizures, structural remodeling, or long-term epileptogenic processes characteristic of chronic epilepsy. Its intended application is therefore complementary to chronic models and focused primarily on mechanistic studies and rapid evaluation of interventions targeting a controlled cortical focus. Second, all experiments were performed under isoflurane anesthesia. Although this provides the stability required for focal microinjection and prolonged multichannel recordings, isoflurane modulates neuronal excitability and may alter seizure dynamics (Becker et al., 2012; Twele et al., 2016; Sorrenti et al., 2021). Surgical manipulation may also induce local inflammatory responses, while small variations in injection depth, 4-AP diffusion, or the relative position of the recording probe and injection site may contribute to inter-animal variability (Balzekas et al., 2016; Tse et al., 2021).

Biological variability represents an additional consideration. Despite standardized experimental procedures, individual susceptibility to 4-AP and pharmacological responsiveness remained variable. Such variability is an inherent feature of experimental neuropharmacology and can persist even under tightly controlled conditions (Löscher, 2024). Both male and female mice were included, although the study was not powered to evaluate sex as an independent experimental factor. Sex and hormonal state may influence seizure susceptibility and antiseizure drug responses, and sex-dependent differences in CBZ pharmacokinetics have been reported (Marino et al., 2012; Christian et al., 2020; Romanescu et al., 2022). Finally, the study was restricted to young adult mice. This age range was selected because preliminary experiments indicated more consistent electrophysiological responses to 4-AP, improving reproducibility, but this controlled selection limits extrapolation to younger or aged populations in which network excitability may differ (Kelly, 2010).

Overall, this study provides a reproducible and experimentally accessible framework for investigating acute focal cortical epileptiform activity. By combining a superficial cortical focus with multichannel electrophysiological recordings and a Python-based multi-biomarker analysis pipeline, the model captures complementary aspects of pathological network activity and their pharmacological sensitivity. This framework may therefore support both mechanistic studies and the quantitative evaluation of pharmacological, optical, and epicortical neuromodulatory strategies.

## DATA AVAILABILITY STATEMENT

The custom Python code used for electrophysiological processing and multi-biomarker analysis will be openly available through the laboratory’s GitHub repository. Repository names, accession numbers, and the corresponding GitHub link will be provided before publication.

## ETHICS STATEMENT

All animal procedures were conducted in accordance with European Union Directive 2010/63/EU and Spanish regulations (Laws 32/2007 and 6/2013 and Real Decreto 53/2013). The experimental procedures were reviewed and approved by the Animal Research Ethics Committee of the University of Barcelona and the Generalitat de Catalunya (license no. 162.24).

## AUTHOR CONTRIBUTIONS

CL-C: Conceptualization, Investigation, Methodology, Formal analysis, Software, Validation, Data curation, Visualization, Writing – original draft, Writing – review & editing. TG: Conceptualization, Supervision, Methodology, Software, Visualization, Writing – review & editing. AP-R: Methodology, Software, Data curation, Writing – review & editing. MVP: Conceptualization, Supervision, Project administration, Resources, Funding acquisition, Visualization, Writing – original draft, Writing – review & editing. All authors contributed to the article and approved the submitted version.

## ACKNOWLEDGEMENTS

The authors thank Sara Hidalgo-Nieves for her excellent technical assistance. Figures 1-4 and 6 were created using BioRender.com.

## GENERATIVE AI STATEMENT

The authors declare that generative AI was used during the preparation of this manuscript. ChatGPT (OpenAI, GPT-5.6 Sol, 2026) was used to assist with language editing, organization, and refinement of the manuscript text. It was not used to generate or analyze experimental data or to create the primary scientific results. All AI-assisted content, including scientific statements, interpretations, and references, was critically reviewed and revised by the authors, who take full responsibility for the accuracy, integrity, and originality of the manuscript.

## SUPPLEMENTARY MATERIAL

The Supplementary Material for this article can be found online at: XX.

## Supplementary Information

**SUPPLEMENTARY FIGURE S1:**
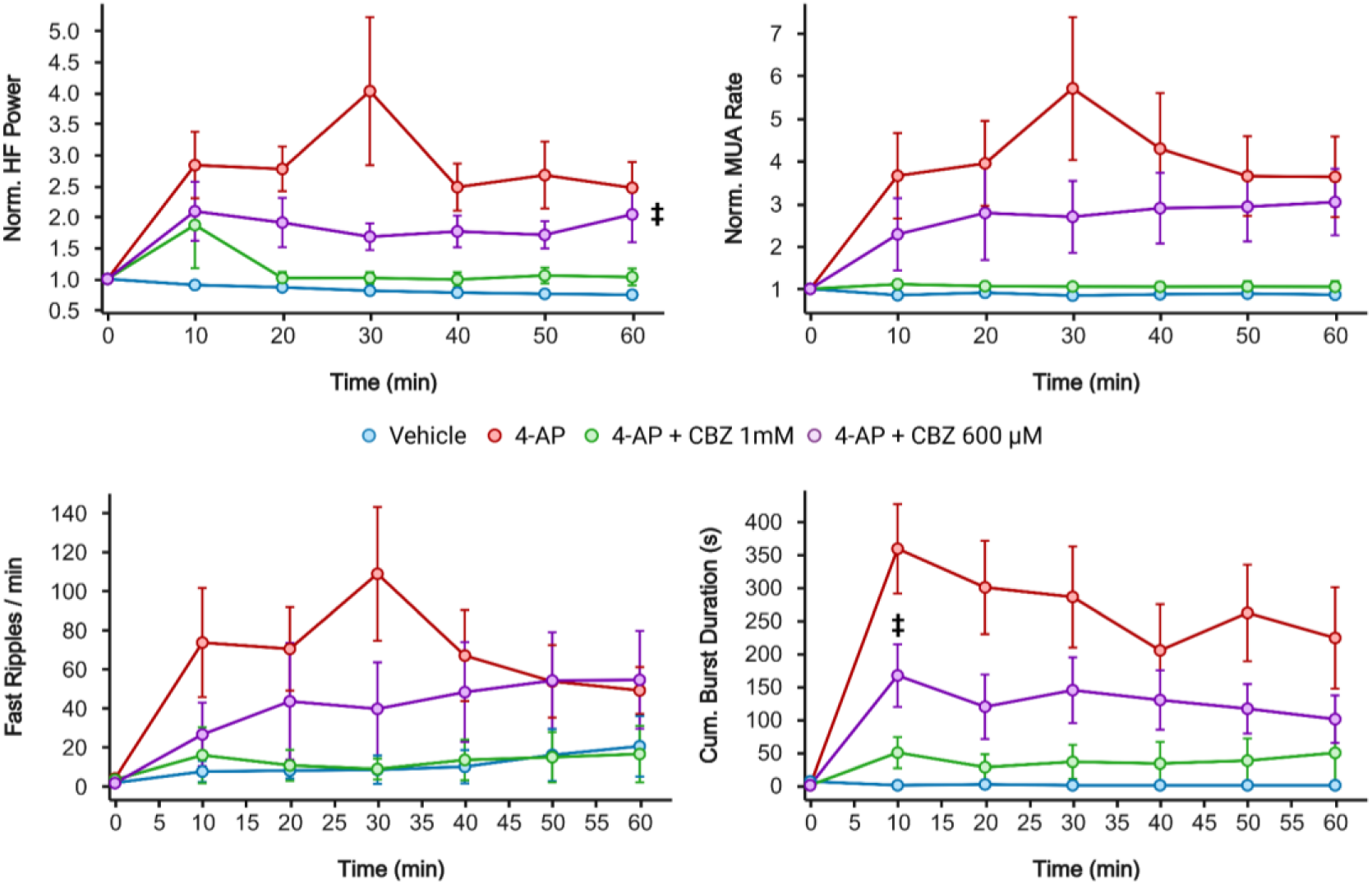
Temporal dynamics of electrophysiological biomarkers including the lower CBZ concentration. Time courses of (**A**) normalized HF power, (**B**) normalized multi-unit activity (MUA), (**C**) fast ripple rate, and (**D**) cumulative burst duration in vehicle, 4-AP (15 mM), 4-AP (15 mM) + CBZ (1 mM), and 4-AP (15 mM) + CBZ (600 μM) groups. HF power and MUA were normalized to each animal’s baseline value (baseline ratio = 1). Baseline is represented at 0 min, followed by six consecutive 10-min post-injection intervals. Data are presented as mean ± SEM. Statistical analyses for this supplementary figure were restricted to the vehicle, 4-AP, and 4-AP + CBZ 600 μM groups; the 1 mM CBZ group is shown for visual reference. Pairwise comparisons were performed following the mixed-design ANOVA with Holm correction for multiple comparisons. Cross symbols indicate significant differences between vehicle and 4-AP + CBZ 600 μM (+p < 0.05).

## REFERENCES

Aanestad, E., Beniczky, S., Olberg, H., and Brogger, J. (2024). Unveiling variability: A systematic review of reproducibility in visual EEG analysis, with focus on seizures. Epileptic Disorders 26, 827–839. doi: 10.1002/epd2.20291

Arida, R. M., Scorza, F. A., de Araujo Peres, C., and Cavalheiro, E. A. (1999). The course of untreated seizures in the pilocarpine model of epilepsy. Epilepsy Research 34, 99–107. doi: 10.1016/S0920-1211(98)00092-8

Assaf, F., and Schiller, Y. (2016). The antiepileptic and ictogenic effects of optogenetic neurostimulation of PV-expressing interneurons. Journal of neurophysiology 116, 1694–1704. doi: 10.1152/jn.00744.2015

Avoli, M., de Curtis, M., Gnatkovsky, V., Gotman, J., Köhling, R., Lévesque, M., et al. (2016). Specific imbalance of excitatory/inhibitory signaling establishes seizure onset pattern in temporal lobe epilepsy. Journal of Neurophysiology 115, 3229–3237. doi: 10.1152/jn.01128.2015

Balzekas, I., Hernandez, J., White, J., and Koh, S. (2016). Confounding effect of EEG implantation surgery: Inadequacy of surgical control in a two hit model of temporal lobe epilepsy. Neuroscience Letters 622, 30–36. doi: 10.1016/j.neulet.2016.04.033

Becker, K., Eder, M., Ranft, A., Von Meyer, L., Zieglgänsberger, W., Kochs, E., et al. (2012). Low dose isoflurane exerts opposing effects on neuronal network excitability in neocortex and hippocampus. PLoS ONE 7, e39346. doi: 10.1371/journal.pone.0039346

Bonaccini-Calia, A., Masvidal-Codina, E., Smith, T. M., Schäfer, N., Rathore, D., Rodríguez-Lucas, E., et al. (2022). Full-bandwidth electrophysiology of seizures and epileptiform activity enabled by flexible graphene microtransistor depth neural probes. Nat Nanotechnol 17, 301–309. doi: 10.1038/s41565-021-01041-9

Bragin, A., Engel, J., Wilson, C. L., Fried, I., and Mathern, G. W. (1999). Hippocampal and entorhinal cortex high-frequency oscillations (100-500 Hz) in human epileptic brain and in kainic acid--treated rats with chronic seizures. Epilepsia 40, 127–137. doi: 10.1111/j.1528-1157.1999.tb02065.x

Camerin, L., Maleeva, G., Gomila-Juaneda, A., Suárez-Pereira, I., Matera, C., Prischich, D., et al. (2024). Photoswitchable carbamazepine analogs for non-invasive neuroinhibition in vivo. *Angew Chem Int Ed Engl*, e202403636. doi: 10.1002/anie.202403636

Christian, C. A., Reddy, D. S., Maguire, J., and Forcelli, P. A. (2020). Sex differences in the epilepsies and associated comorbidities: Implications for use and development of pharmacotherapies. Pharmacological Reviews 72, 767–800. doi: 10.1124/pr.119.017392

Delgado-Sallent, C., Gener, T., Nebot, P., López-Cabezón, C., and Puig, M. V. (2023). Neural substrates of cognitive impairment in a NMDAR hypofunction mouse model of schizophrenia and partial rescue by risperidone. Frontiers in Cellular Neuroscience 17. doi: 10.3389/fncel.2023.1152248

Esteller, R., Echauz, J., Tcheng, T., Litt, B., and Pless, B. (2001). Line length: an efficient feature for seizure onset detection., in Conference of the IEEE Engineering in Medicine and Biology Society, 1707–1710 vol.2. doi: 10.1109/IEMBS.2001.1020545

Hamani, C., and Mello, L. E. A. M. (2002). Spontaneous recurrent seizures and neuropathology in the chronic phase of the pilocarpine and picrotoxin model epilepsy. Neurological Research 24, 199–209. doi: 10.1179/016164102101199611

Heining, K., Kilias, A., Janz, P., Häussler, U., Kumar, A., Haas, C. A., et al. (2019). Bursts with high and low load of epileptiform spikes show context-dependent correlations in epileptic mice. eNeuro 6, ENEURO.0299-18.2019. doi: 10.1523/ENEURO.0299-18.2019

Inouye, T., Shinosaki, K., Sakamoto, H., Toi, S., Ukai, S., Iyama, A., et al. (1991). Quantification of EEG irregularity by use of the entropy of the power spectrum. Electroencephalogr Clin Neurophysiol 79, 204–210. doi: 10.1016/0013-4694(91)90138-t

Jia, F., Yue, M., Chandra, D., Homanics, G. E., Goldstein, P. A., and Harrison, N. L. (2008). Isoflurane is a potent modulator of extrasynaptic GABA(A) receptors in the thalamus. J Pharmacol Exp Ther 324, 1127–1135. doi: 10.1124/jpet.107.134569

Jiang, M., and Wang, Y. (2025). Refinement of the pilocarpine-induced status epilepticus model in mice to improve mortality outcomes. Front. Neurosci. 19. doi: 10.3389/fnins.2025.1592014

Jing, J., Ge, W., Struck, A. F., Fernandes, M. B., Hong, S., An, S., et al. (2023). Interrater reliability of expert electroencephalographers identifying seizures and rhythmic and periodic patterns in EEGs. Neurology 100, e1737–e1749. doi: 10.1212/WNL.0000000000201670

Kamintsky, L., van Hameren, G., Weissberg, I., Moradi, P., Prager, O., Ahmad, A. A., et al. (2025). An algorithm for seizure detection in rodents. Epilepsia Open. doi: 10.1002/epi4.70070

Kandratavicius, L., Balista, P. A., Lopes-Aguiar, C., Ruggiero, R. N., Umeoka, E. H., Garcia-Cairasco, N., et al. (2014). Animal models of epilepsy: use and limitations. NDT 10, 1693–1705. doi: 10.2147/NDT.S50371

Kelly, K. M. (2010). Aging models of acute seizures and epilepsy. Epilepsy Curr 10, 15–20. doi: 10.1111/j.1535-7511.2009.01341.x

Kuo, C. C., Chen, R. S., Lu, L., and Chen, R. C. (1997). Carbamazepine inhibition of neuronal Na+ currents: quantitative distinction from phenytoin and possible therapeutic implications. Mol Pharmacol 51, 1077–1083. doi: 10.1124/mol.51.6.1077

Lévesque, M., and Avoli, M. (2013). The kainic acid model of temporal lobe epilepsy. Neuroscience & Biobehavioral Reviews 37, 2887–2899. doi: 10.1016/j.neubiorev.2013.10.011

Löscher, W. (2017). Animal models of seizures and epilepsy: Past, present, and future role for the discovery of antiseizure drugs. Neurochem Res 42, 1873–1888. doi: 10.1007/s11064-017-2222-z

Löscher, W. (2024). Of mice and men: The inter-individual variability of the brain’s response to drugs. eNeuro 11, ENEURO.0518-23.2024. doi: 10.1523/ENEURO.0518-23.2024

Marino, S. E., Birnbaum, A. K., Leppik, I. E., Conway, J. M., Musib, L. C., Brundage, R. C., et al. (2012). Steady-state carbamazepine pharmacokinetics following oral and stable-labeled intravenous administration in epilepsy patients: Effects of race and sex. Clin Pharmacol Ther 91, 483–488. doi: 10.1038/clpt.2011.251

Nagappan, S., Liu, L., Fetcho, R., Nguyen, J., Nishimura, N., Radwanski, R. E., et al. (2019). In vivo femtosecond laser subsurface cortical microtransections attenuate acute rat focal seizures. Cereb Cortex 29, 3415–3426. doi: 10.1093/cercor/bhy210

Prokop, M., Esparza-Iaizzo, M., Masvidal-Codina, E., Illa, X., Codadu, N. K., Rathore, D., et al. (2026). An artefact-resilient wide bandwidth bidirectional graphene neural interface. Nat Commun 17, 6536. doi: 10.1038/s41467-026-73790-x

Quiroga, R. Q., Nadasdy, Z., and Ben-Shaul, Y. (2004). Unsupervised spike detection and sorting with wavelets and superparamagnetic clustering. Neural Comput. 16, 1661–1687. doi: 10.1162/089976604774201631

Romanescu, M., Buda, V., Lombrea, A., Andor, M., Ledeti, I., Suciu, M., et al. (2022). Sex-related differences in pharmacological response to CNS drugs: A narrative review. JPM 12, 907. doi: 10.3390/jpm12060907

Sorrenti, V., Cecchetto, C., Maschietto, M., Fortinguerra, S., Buriani, A., and Vassanelli, S. (2021). Understanding the effects of anesthesia on cortical electrophysiological recordings: A scoping review. IJMS 22, 1286. doi: 10.3390/ijms22031286

Staba, R. J., Wilson, C. L., Bragin, A., Fried, I., and Engel, J. (2002). Quantitative analysis of high-frequency oscillations (80–500 Hz) recorded in human epileptic hippocampus and entorhinal cortex. Journal of Neurophysiology 88, 1743–1752. doi: 10.1152/jn.2002.88.4.1743

Thijs, R. D., Surges, R., O’Brien, T. J., and Sander, J. W. (2019). Epilepsy in adults. Lancet 393, 689–701. doi: 10.1016/S0140-6736(18)32596-0

Tse, K., Beamer, E., Simpson, D., Beynon, R. J., Sills, G. J., and Thippeswamy, T. (2021). The impacts of surgery and intracerebral electrodes in C57BL/6J mouse kainate model of epileptogenesis: Seizure threshold, proteomics, and cytokine profiles. Front. Neurol. 12. doi: 10.3389/fneur.2021.625017

Twele, F., Schidlitzki, A., Töllner, K., and Löscher, W. (2017). The intrahippocampal kainate mouse model of mesial temporal lobe epilepsy: Lack of electrographic seizure-like events in sham controls. Epilepsia Open 2, 180–187. doi: 10.1002/epi4.12044

Twele, F., Töllner, K., Brandt, C., and Löscher, W. (2016). Significant effects of sex, strain, and anesthesia in the intrahippocampal kainate mouse model of mesial temporal lobe epilepsy. Epilepsy Behav 55, 47–56. doi: 10.1016/j.yebeh.2015.11.027

Wang, S. H., Ferrari, P., Arnulfo, G., Marzulli, M., Degras, D., Myrov, V., et al. (2026). Toward unified biomarkers for focal epilepsy. J Neurosci 46, e0879252025. doi: 10.1523/JNEUROSCI.0879-25.2025

Weiss, S. A., Song, I., Leng, M., Pastore, T., Slezak, D., Waldman, Z., et al. (2020). Ripples have distinct spectral properties and phase-amplitude coupling with slow waves, but indistinct unit firing, in human epileptogenic hippocampus. Front Neurol 11, 174. doi: 10.3389/fneur.2020.00174

Yang, F., Li, J., Song, Y., Zhao, M., Niemeyer, J. E., Luo, P., et al. (2021). Mesoscopic mapping of ictal neurovascular coupling in awake behaving mice using optical spectroscopy and genetically encoded calcium indicators. Front Neurosci 15, 704834. doi: 10.3389/fnins.2021.704834

Zhao, M., McGarry, L. M., Ma, H., Harris, S., Berwick, J., Yuste, R., et al. (2015). Optical triggered seizures using a caged 4-Aminopyridine. Front. Neurosci. 9. doi: 10.3389/fnins.2015.00025

Zuev, V. A., Salmagambetova, E. G., Djakov, S. N., and Utkin, L. V. (2025). Automated video-EEG analysis in epilepsy studies: A narrative review of advances and challenges. J Med Syst 49, 142. doi: 10.1007/s10916-025-02274-0

